# mirtronDB: a mirtron knowledge base

**DOI:** 10.1101/429522

**Authors:** Bruno Henrique Ribeiro Da Fonseca, Douglas Silva Domingues, Alexandre Rossi Paschoal

## Abstract

**Motivation:** Mirtrons are originated from short introns with atypical cleavage from the miRNA canonical pathway by using the splicing mechanism. Several studies describe mirtrons in chordates, invertebrates and plants but in the current literature there is no repository that centralizes and organizes these public and available data. To fill this gap, we created the first knowledge database dedicated to mirtron, called mirtronDB, available at http://mirtrondb.cp.utfpr.edu.br/. MirtronDB has a total of 1,407 mirtron precursors and 2,426 mirtron mature sequences in 18 species.

**Results:** Through a user-friendly interface, users can browse and search mirtrons by organism, organism group, type and name. MirtronDB is a specialized resource to explore mirtrons and their regulations, providing free, user-friendly access to knowledge on mirtron data.

**Availability:** MirtronDB is available at http://mirtrondb.cp.utfpr.edu.br/.

**Supplementary information:** Supplementary data are available.

## 1 Introduction

Studies in model organisms identified that some short hairpin introns presented similar characteristics to miRNAs, however using the splicing mechanism as the first stage of the miRNA biogenesis cleavage (Ruby, J. et al., 2007). These non-canonical miRNAs described in small introns are collectively called “mirtrons” (Okamura, K. et al., 2007).

Mirtron deregulation was later identified as potential source of several human pathologies (Qu, Z. and Adelson, D., 2012) and its use in therapeutic treatments has been considered (Curtis, H. et al., 2017). In plants, an intronic miRNA, miR838, has been located on the DCL1 primary transcript in *A. thaliana*, suggesting a feedback loop for the autoregulation of miRNA biogenesis (Rajagopalan, R. et al., 2006 and Budak, H. and Akpinar, B., 2015).

In the literature, there is no repository for accessing knowledge on mirtron data. Not even miRBase (Griffiths-Jones, S. et al., 2006), the miRNA state-of-the-art repository, has specific analysis for mirtrons. Until November 2017, we identified 22 articles that have available public mirtron data. However, these datasets are dispersed, with no standardization or organization.

Organization of these data will enable studies on mirtron characteristics, roles, and interactions in organisms, among other potential scientific advances. In this context, to fill this gap, we provide mirtronDB (http://mirtrondb.cp.utfpr.edu.br/), a central mirtron knowledge data repository. For that, we modelled a total of 1,407 mirtron precursors and 2,426 mature mirtrons from 18 species (chordates, invertebrates and plants) based on published available literature.

MirtronDB has an online user-friendly interface for the user to search, browser, visualize, and download information about mirtrons. All datasets are publicly available in several formats. The user has access to: (i) precursor mirtron similarity analysis; (ii) target gene predictions; and (iii) ceRNA predictions in plants. We expect this resource can increase the amount studies on mirtrons.

## 2 Materials and methods

MirtronDB was built in four steps: (1) Data collection; (2) Data modelling; (3) Data analysis; and (4) Website interface (Supplementary Figure S1).

### 2.1 Mirtrons data collection

We collected the mirtron data available from June 2007 to November 2017 searching by the term “mirtron OR mirtrons” in the fields title/abstract in NCBI PubMed (Supplemental Table S1) and in papers cited in them. The articles selected were manually analyzed and redundancies were removed. We created a standardized name: “organism name abbreviation + the word ‘mirtron’ + ID, and for mature we add the arm”. We built a database (Supplementary Figure S2) and automatically imported the information.

### 2.2 Data analysis

#### 2.2.1 Similarity analysis among organisms

We extracted genomic information from several sources (Supplementary Table S2). We performed a BLASTN alignment, version 2.6 (Camacho, C. et al., 2009), between all precursor mirtrons against all other species genomes. We retained results that have above 95% query coverage and identity.

#### 2.2.2 Mirtrons and miRNAs similarity analysis

The mature mirtrons were aligned to all miRNAs from miRBase v22 (Griffiths-Jones, S. et al., 2006) using the CD-HIT-EST-2D tool (Huang, B. et al., 2010) and considering the alignment of 9 nucleotides (nt) at 0.98 of cutoff identity.

#### 2.2.3 Target gene prediction

We predicted the targets gene for *H. sapiens* and plants. For human, we used TargetScan (Agarwal, V. et al., 2015) with default parameters and psRNATarget (Dai, X. et al., 2017) was used in plants (seed region parameter from 2 to 8 nt).

#### 2.2.4 ceRNA prediction in plants

We used TAPIR (Bonnet, E. et al., 2010, version 1.2) to predict ceRNA in plants, with default parameters. All mature mirtrons were compared against all lncRNAs from GreeNC database (Gallart, A. et al., 2015, version 1.12).

#### 2.2.5 Website implementation

MirtronDB was developed using HTML 5, PHP 7.0, CSS 4.0, and Bootstrap 3.3. The network visualization was done using Cytoscape.js (Franz, M. et al., 2015) and PostgreSQL as relational database system.

## 3 Results

### 3.1 mirtronDB: database content

We found a total of 1,407 precursor mirtrons and 2,426 mature mirtrons in 18 species and we extracted functional information, when available. All mirtrons collected are detailed in Supplementary Table S3. The Supplementary Figure S3 presents the cumulative distribution of mirtron information per year.

### 3.2 Precursor mirtron similarity analysis

We obtained 944 aligned precursor mitrons, where 896 were aligned in chordates (94.9%), 46 in invertebrates (4.9%) and 2 in plants (0.2%) (Supplementary Table S4). Only four species had more than 3 mirtrons aligned in another genome: *H. sapiens*, *M. mulatta*, *P. troglodytes* and *D. melanogaster*.

### 3.3 Mature mirtron characterization

In chordates and invertebrates, most mature mirtrons have 22 nt (32.1%) and, in plants, most mature mirtrons (28%) have 21 nt (Supplementary Table S5 and Figure S4). We obtained logo sequences for mirtron arms where chordates present more GC bases than invertebrates and plants (Supplementary Figure S5).

### 3.4 Mirtrons availability in miRBase

We investigated if mature mirtron sequences were represented in the miRBase. We observed that 966 mirtrons (39.8%) are available in the miRBase, reinforcing the novelty provided by mirtronDB (Supplementary Table S6).

### 3.5 Target gene and ceRNA analysis

We identified a total of 512,298 and 3,884 potential targets gene predictions in humans and plants, respectively (Supplemental Table S3). In plants, we also verified if mirtrons could act as ceRNA candidates, where a total of 1,738 potential interactions were found (Supplementary Table S7).

### 3.6 mirtronDB: user interfaces

MirtronDB portal provides a user-friendly web interface, various query functions, and graphical visualization to access mirtron knowledge. We highlight that in “Search” the users can query mirtrons by organism, organism group, type and mirtron name, in “Network” the users can build mirtrons/target-specific networks, and the analysis results are presented per mirtron in query result.

## 4 Discussion

MirtronDB is a database that standardizes, integrates and provides mirtron data available from literature. We highlight that (i) all data collected is available in several formats; (ii) curated data make this repository a mirtron information reference; (iii) sequence, structure and conservation analysis are provided; and (iv) potential targets and ceRNA in mirtrons are also investigated.

Data availability facilitates the development of new studies in biology. For example, the similarity analysis and target gene prediction could then be tested in a wet lab. MirtronDB data could also provide novel standardized approaches to analyze mirtrons in organisms that do not have them described yet.

## 5 Conclusion

MirtronDB is a comprehensive database about mirtrons that allows users to query mirtron data and download them. The analyses presented in this paper provides initial mirtron characterization and can be used as a guide about mirtrons' potential. This repository also has the potential to promote advances in computational biology, since once consolidated data available can now be used to improve or model mirtron approaches and biological experiments.

